# PrePPI: A structure informed proteome-wide database of protein-protein interactions

**DOI:** 10.1101/2023.02.27.530276

**Authors:** Donald Petrey, Haiqing Zhao, Stephen Trudeau, Diana Murray, Barry Honig

**Affiliations:** Department of Systems Biology, Columbia University Irving Medical Center, New York, NY 10032, USA; Department of Biochemistry and Molecular Biophysics, Columbia University Irving Medical Center, New York, NY 10032, USA; Department of Medicine, Columbia University, New York, NY 10032, USA; Zuckerman Mind Brain and Behavior Institute, Columbia University, New York, NY, 10027, USA

**Keywords:** Protein-protein interactions, database, AlphaFold models, structural modeling, non-structural clues

## Abstract

We present an updated version of the Predicting Protein-Protein Interactions (PrePPI) webserver which predicts PPIs on a proteome-wide scale. PrePPI combines structural and non-structural clues within a Bayesian framework to compute a likelihood ratio (LR) for essentially every possible pair of proteins in a proteome; the current database is for the human interactome. The structural modeling (SM) clue is derived from template-based modeling and its application on a proteome-wide scale is enabled by a unique scoring function used to evaluate a putative complex. The updated version of PrePPI leverages AlphaFold structures that are parsed into individual domains. As has been demonstrated in earlier applications, PrePPI performs extremely well as measured by receiver operating characteristic curves derived from testing on *E. coli* and human protein-protein interaction (PPI) databases. A PrePPI database of ~1.3 million human PPIs can be queried with a webserver application that comprises multiple functionalities for examining query proteins, template complexes, 3D models for predicted complexes, and related features (https://honiglab.c2b2.columbia.edu/PrePPI). PrePPI is a state-of-the-art resource that offers an unprecedented structure-informed view of the human interactome.

**Graphic Abstract:** 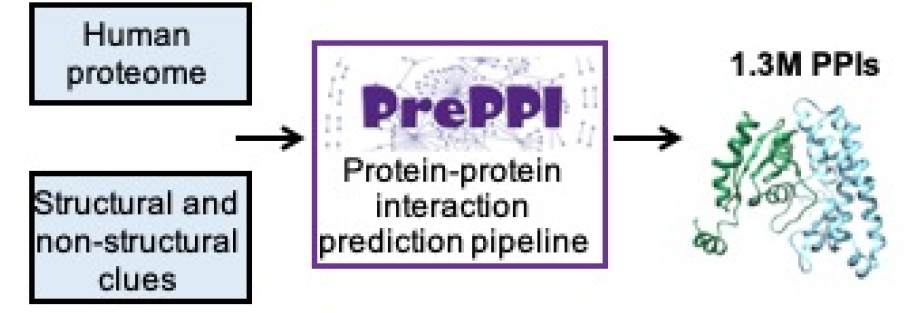

## Introduction

The identification of proteins that interact with one another is a challenging problem of central importance in fundamental biology and in medicine. Protein-protein interactions (PPIs) is a widely used term which has multiple meanings. Two proteins can interact with one another directly either by forming a binary physical complex or by being in physical contact in the context of a multi-protein complex. Indirect interactions can include two proteins that are part of a complex, but are not in physical contact, or that are part of a pathway or network that mediates their interaction. Multiple experimental and computational tools are available to detect or predict PPIs, and their results are compiled in multiple databases. Here we report a new version of our Predicting Protein-Protein Interactions (PrePPI) database [1, 2], describe its unique features, and compare its performance to that of other databases. We also place PrePPI’s prediction algorithm in the context of recent structure-based, co-evolution, and deep learning-based developments in the prediction of PPIs.

The key element of the PrePPI algorithm, which is summarized in Figure 1, is proteome-wide template-based modeling of PPIs, both direct and indirect. Not accounting for splice variants and posttranslational modifications, there are ~200 million possible pairwise combinations of human proteins. However, since we consider full proteins as well as their individual domains, we need to examine ~4.55 billion pairwise interactions and, since we make multiple interaction models for each pair, the number of pairwise combinations evaluated is in the 10s of billions (see Methods). PrePPI’s ability to consider such a large number of potential PPIs is enabled by an efficient scoring function which is based on the similarity of the modeled interface to the interface of a known complex in the Protein Data Bank (PDB) [3]. We highlight these points because it is important to distinguish our goals from standard template-based modeling. Further, we are not necessarily trying to produce an accurate model of the complex as might be judged, for example, in the CAPRI (Critical Assessment of PRediction of Interactions) experiment [4] – although obviously a better model will produce a more reliable prediction. Rather, our hypothesis is that, in the derivation of a structural modeling score, our models are good enough to provide a clue that two proteins form a physical complex. Thus, a model that would score poorly according to CAPRI metrics might be reliable enough to provide a yes or no prediction as to whether two proteins interact and, in addition, produce a low-resolution structural pose for the interaction. As discussed below, PrePPI uses non-structural information as well. For example, if two proteins are co-expressed and have a good structural modeling (SM) score, the likelihood of an interaction, as given in PrePPI by a naïve Bayesian network, will increase. A PPI with low SM score but high non-structural score suggests that the interaction is indirect.

**Figure 1:**
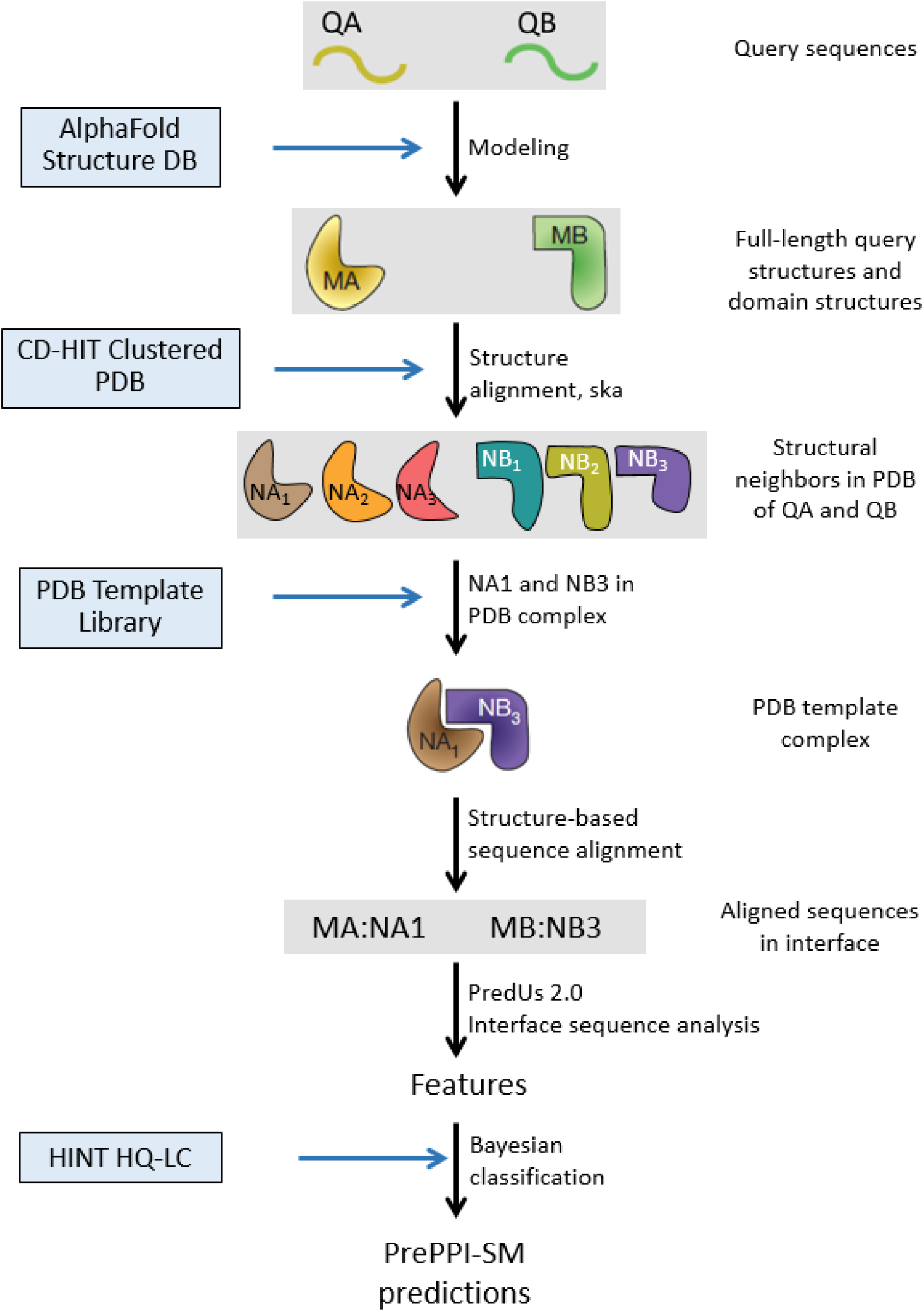
PrePPI’s structural modeling (SM) pipeline. Structures for query proteins, QA and QB, are taken from the AlphaFold Protein Structure Database [13] and parsed into domains with definitions from the Conserved Domain Database (CDD) [22]. Structural neighbors in the PDB [3] for full length protein and domain structures are obtained from the ska structural alignment program [31]. If structural neighbors of two query proteins appear together in a PDB complex, this structure defines a template, NA1:NB3, used to create a structure-based sequence alignment with which an interface for the query proteins, QA:QB, is evaluated based on the overlap of the query and template residues [1]. The interaction is then scored based on a number of features [1, 2] and trained on the HINT HQ-LC database [10], as the positive set, and a negative set described in Methods to produce a fully connected Bayesian network used to evaluate the model.

Testing and validating computational predictions is a complicated challenge since experimental databases themselves contain sources of uncertainty and the degree of overlap between them is still quite low in spite of the proliferation of observations from high-throughput screens. Moreover, they are often based on different definitions of PPIs. Mass spectrometry-derived databases (e.g. Bioplex 3.0 [5]) focus explicitly on multi-protein complexes [6] while Y2H-based databases (e.g. HuRI [7]) focus on binary interactions. Among derived databases, the widely used STRING database [8] has a category for physical interactions but does not distinguish binary interactions from those in multi-protein complexes whereas databases such as APID [9] and HINT [10] include both direct and indirect interactions. As depicted in Table 1, overlap between these various databases is limited (see Methods for a description of each database). Of note, Interactome 3D which contains PDB structures and high quality homology models is well represented in most of the databases but, the HINT high-quality literature-curated database (HINT HQ-LC) contains the highest percentage of Interactome3D structures.

**Table 1.**
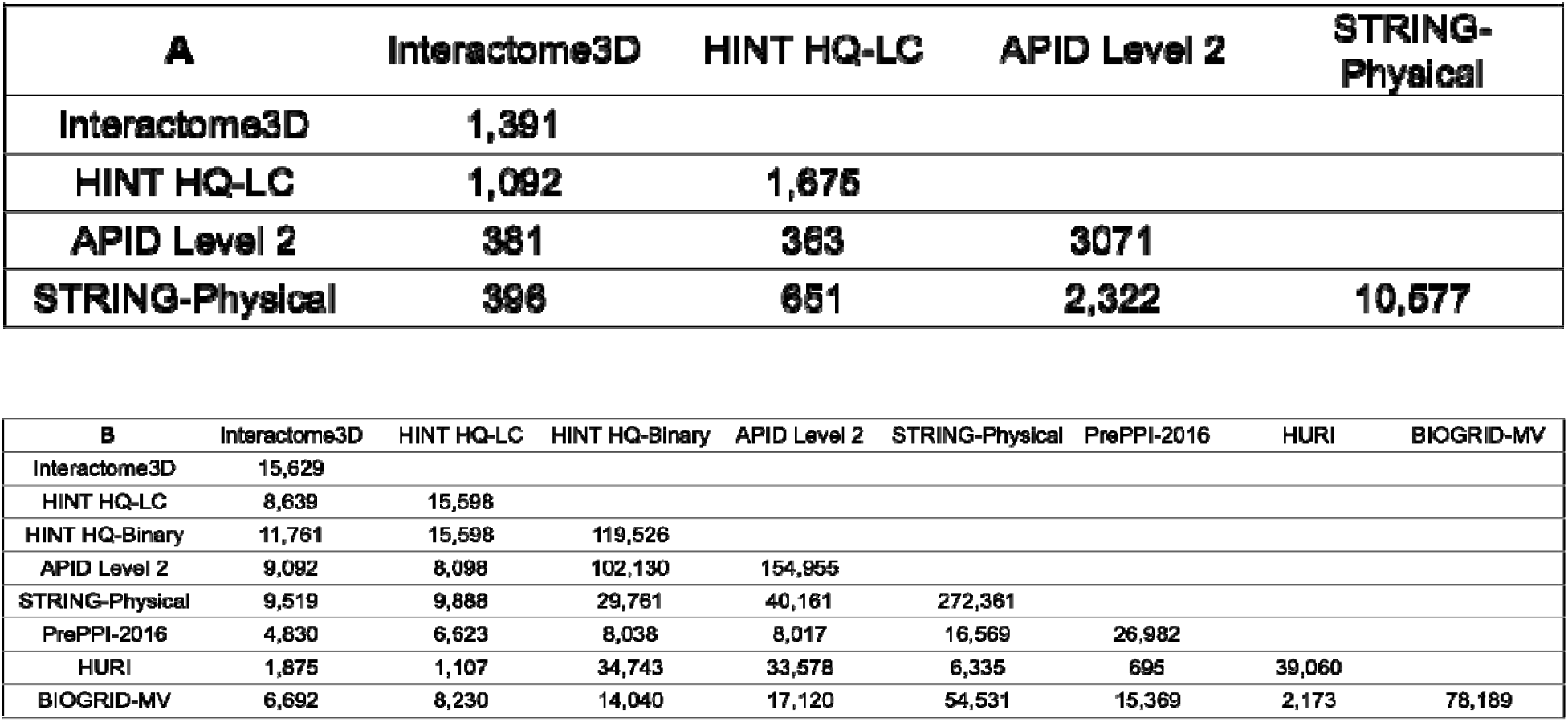
Overlap among PPI databases: The number of overlapping entries between Hint HQ-LC, Interactome3D, APID Level2 and STRING-Physical are listed for **A**. *E. coli* and **B**. Human. The numbers for Interactome 3D include PDB structures and easily constructed homology models.

In earlier versions of PrePPI [1, 2], training was done on yeast PPIs and testing was done on human interactions, with the true positive dataset comprising PPIs with at least two literature references. No attempt was made at the time to train on datasets of binary physical interactions since PrePPI predicts both direct and indirect interactions. Here we have taken a more refined approach, training the structural modeling component of PrePPI on HINT HQ-LC [10].

In order to evaluate PrePPI’s structure-based algorithm, we have used *Escherichia coli* K-12 (here *E. coli*) as a test organism and compared predictions from PrePPI’s structural modeling component to predictions from the threading component of Threpp [11]. Technology closely related to Threpp powers the PEPPI server [12] which, like PrePPI, uses Bayesian statistics to integrate structural and non-structural information. But in contrast to the PrePPI, the PEPPI webserver allows a user to input only two protein sequences at a time while, as described below, the PrePPI database of human PPIs contains about 200 million entries with the highest confidence predictions (~1.3M) appearing in the online application that can be queried in multiple ways including, for example, inputting a single protein and outputting all predicted binding partners.

Compared to previous versions of PrePPI, in addition to improved training, features of the current version include the replacement of homology models with models from the AlphaFold Protein Structure Database [13] leading to increased structural coverage of the proteome, separate training of the structural modeling and non-structural components, a refined definition of PDB template complexes [3], the implementation of a more accurate algorithm PredUs 2.0 for predicting interfacial residues [14], and a website with expanded functionality. We believe that PrePPI is a unique resource that generates novel hypotheses for the existence of PPIs, both direct and indirect. Moreover, given the ongoing developments in the use of deep learning-based approaches to predict the structure of binary complexes, PrePPI predictions can be used as a starting point for the construction of accurate structural models.

## Results

### Testing on experimental databases

#### E. coli

We have chosen to test the SM score on *E. coli*, in part for comparison with Threpp [11] and in part to assess the applicability of our human-trained Bayesian network (see below) to another organism. PrePPI for *E. coli* was trained on human HINT HQ-LC [10] (see Methods). Table 2A presents area under the ROC curve (AUROC) values for the structural modeling component of PrePPI (PrePPI-SM) and the threading component of Threpp (Threpp-Threading) [11, 12] for *E. coli* evaluated on three datasets: *E. coli* HINT HQ-LC, Interactome3D, and GS-Threpp [15], the gold standard data set of 763 PPIs on which Threpp was previously tested [11]. Both methods yield good results when tested on HINT HQ-LC (AUROC values 0.88 and 0.81 for PrePPI-SM and Threpp-Threading, respectively) and Interactome3D (AUROC values 0.95 and 0.85) but performance degrades (AUROC values 0.67 and 0.65) on GS-Threpp. PrePPI-SM performs quite well on HINT HQ-LC on which it was trained and performance improves on Interactome3D which is comprised effectively of PDB complexes or close homologs [16]. As can be seen in Table 1A, HINT HQ-LC has a large intersection with Interactome3D (65%). The slight difference in performance may arise if some of the interactions in HINT HQ-LC are not readily homology-modeled. Overall, the PrePPI-SM results are somewhat better than those obtained with Threpp-Threading but it is reassuring that two different structure-based methods yield very similar performance and, in particular, that a proteome-wide method such as PrePPI is of comparable accuracy to a method that uses a more complex and computationally intensive scoring function to evaluate structural models.

**Table 2.** Area Under Roc curve, AUROC, for different test sets. **A.** *E coli*. The structural modeling performance of PrePPI compared to that of Threpp-Threading, both tested on Interactome3D, Hint HQ-LC and GS-Threpp. **B.** *Human*. The performance of PrePPI-SM and PrePPI-total tested on Hint HQ-LC and the PrePPI 2016 high confidence set.

| **A** | **HINT HQ-LC** | **Interactome3D** | **GS-Threpp** |
| --- | --- | --- | --- |
| **PrePPI-SM** | **0.88** | **0.95** | **0.67** |
| **Threpp-Threading** | **0.81** | **0.85** | **0.65** |

**Table 2.** Area Under Roc curve, AUROC, for different test sets. **A. E. coli**. The structural modeling performance of PrePPI compared to that of Threpp-Threading, both tested on
| **B** | **HINT HQ-LC** | **PrePPI-2016** |
| --- | --- | --- |
| **PrePPI-SM** | **0.83** | **0.73** |
| **PrePPI-Total** | **0.77** | **0.89** |

#### Human

Table 2B presents AUROC values for PrePPI-SM and PrePPI-Total, where the latter corresponds to the predicted score with all sources of evidence (Figure 1), for testing on HINT HQ-LC and the high confidence (HC) set we assembled in 2016, PrePPI-2016 [2]. PrePPI-SM performs very well on HINT HQ-LC (AUC = 0.83) but performance degrades on PrePPI-2016 (AUC = 0.73). We attribute the difference to the fact that HINT HQ-LC was designed to encompass experimentally observed direct PPIs and, thus, has significant overlap (56%) with Interactome3D [16] (Table 1B) while PrePPI-2016 contains many indirect interactions (19% overlap with Interactome3D). Consistent with this explanation, the difference in performance between the use of just structural clues or the combination of structural and non-structural clues for testing on HINT HQ-LC (AUROC = 0.83 for PrePPI-SM and 0.77 for PrePPI-Total) is small, whereas the AUROC for testing on the 2016 HC set decreases from 0.89 for PrePPI-Total to 0.73 for PrePPI-SM, indicating that PrePPI-Total successfully captures both structural and non-structural evidence.

Table S1 contains AUROC values for PrePPI-Total tested on a number of PPI databases. The values vary over a wide range which appears to reflect underlying differences in the databases as delineated in Table 1. As summarized in Methods, HURI [7], HINT HQ-Binary [10] and APID Level 2 [9] contain many Y2H results, STRING-Physical [17] contains many direct and indirect physical interactions while BioGRID-MV [18] infers PPIs from a large set of experimental methods. HINT HQ-LC is derived from binary interactions that have at least two literature references and, in that sense, is most closely related to PrePPI-2016. Agreement between PrePPI and HURI is quite limited (see Luck et al. [7]) for a discussion of HURI’s overlap with other databases). Of course, it is impossible to know how many predicted PPIs that do not appear in any database are actually true positives. Indeed PrePPI’s goal is to discover PPIs that do not appear in known databases. Based on experimental tests and applications summarized in the Discussion, PrePPI has already proved to be a reliable source of novel PPIs.

To place PrePPI predictions in the context of deep learning approaches, we compared PrePPI performance to that of D-SCRIPT [19], a proteome-wide method for predicting physical interactions between two proteins given just their sequences. Similar to PrePPI, D-SCRIPT was trained on human PPIs and then predicts PPIs for both human and *E. coli*, but training and testing were performed with PPIs from the STRING database [17] whereas PrePPI used HINT HQ-LC [10] (see comparisons in Table 1A and B). In spite of the differences in training and testing sets, the performance, as judged by AUROC values, is similar for both *E. coli* (PrePPI-SM: 0.88, D-SCRIPT: 0.86) and human (PrePPI-SM: 0.83, D-SCRIPT: 0.83) PPIs. Given the low overlap between the HINT HQ-LC and STRING-Physical databases, the strong performance of both methods suggests they are highly complementary, not only in methodological terms but also in the type of information they encompass.

#### The PrePPI database

The full PrePPI database contains predictions for ~200 million PPIs. Even though interaction models are evaluated for a protein and its constituent domains, only the highest scoring interaction for a given protein pair is included in the database. Hence, the set of 200 million PPIs corresponds to near total coverage of all possible interactions among 20K proteins. The online database contains about 1.3M human PPIs of which about 370K are likely direct physical interactons. PPIs that appear in the online database either are associated with an FPR < 0.005 (LR > 379) or have the maximum value of LR(SM) or LR(protein-peptide) > 100. Our experience has been that interactions that meet this latter criterion constitute high-confidence physical interactions and, indeed, are associated with an FPR < 0.001 when tested, with 10-fold cross-validation, on the structure-rich HINT HQ-LC database.

##### PrePPI website (https://honiglab.c2b2.columbia.edu/PrePPI/)

When a user inputs a UniProt ID or gene name for a query protein, the website returns several features of the protein and its interactors: 1) the names and functional information for the query protein derived from UniProt; 2) the sequence of the full-length query protein as well as its domains, all of which can be viewed in a protein-centric structure viewer; 3) a list of PrePPI-predicted interactors of the query protein and associated scores for the features incorporated in the PrePPI algorithm, and, if they exist for a given PPI, links to external databases that compile interactions based on experiments and literature; 4) an interaction-centric structure viewer that shows the 3D model for a given PPI and, depending on selections by the user, the template PDB complex and the structure superposition of the query structures on the template (Figure 1); 5) functional annotations for the query protein, derived from gene set enrichment analysis of the protein’s interactors ranked according to the PrePPI-Total score [2]; 6) annotations of the full-length query protein sequence for disordered regions [20]; and 7) annotations of the full-length query protein sequence for interfacial residues as predicted by PredUs 2.0 [14] that is used in the PrePPI SM scoring function (Figure 1).

## Discussion

The PrePPI database was first reported in 2012 [1] and updated in 2016 [2]. Its unique features include a fast structure-based scoring function that enables proteome-wide protein-protein interface evaluation and the integration of structural and non-structural clues for an interaction. The current version of PrePPI has been improved in a number of ways: 1) Most notably, our in-house homology model database has been replaced with structures from the AlphaFold Protein Structure Database [13] for individual proteins and their domains as annotated by the Conserved Domain Database (CDD) [21]. As explained in Methods, use of the AF/CDD database requires the scoring 10s of billions of interaction models. This scoring takes about a day using ~2000 CPU processors. 2) The training of structure-based versus non-structural clues is performed separately. Specifically, the structure-informed predictions are trained with the HINT HQ-LC database [10] while non-structural features are derived as implemented previously [2] and trained on databases with a predominance of non-structural information. 3) The method to extract non-crystallographic protein-protein interfaces from the PDB has been revised. 4) A more accurate algorithm, PredUs 2.0, was implemented for predicting interfacial residues on protein surfaces [14]. 5) New website features are as described above.

We are not aware of any structure-informed database comparable in scope to PrePPI. Many of its predictions have not been previously observed since use of 3D structure information, especially in matching protein structures to PPI template complexes from the PDB, identifies many interactions that would be undetectable with sequence-based methods. PrePPI performance is comparable to that of high-throughput experimental methods [1, 2]. Moreover, experimental validation has already confirmed the reliability of many novel predictions: 1) In the original PrePPI paper [1], 17 out of 21 predictions were confirmed with co-IP assays; 2) In our study of virus/human interactions with the P-HIPSTer database, which is based on the PrePPI pipeline [22], PrePPI predictions yielded a 76% precision as judged by co-IP experiments; 3) PrePPI is a central feature in the OncoSig algorithm that generated a lung cancer adenocarcinoma (LUAD) signaling PPI network for KRAS that recapitulated published KRAS biology and identified novel proteins synthetic lethal with an oncogenic mutated form of KRAS that is constitutively activated, 18 of 21 of which were validated in 3D spheroid models for LUAD [23]. Thus, based on results in a wide range of contexts, PrePPI predictions are associated with a precision of ~75-80%.

Of course, not all PrePPI predictions are correct but, as highlighted in the previous paragraph, they appear sufficiently accurate to generate hypotheses that drive biological discovery. Moreover, for direct binary PPIs, a model that appears in the database can be used as a basis for lower throughput approaches such as proteinprotein docking or deep learning algorithms such as AlphaFold multimer [24] which likely generate models that are more accurate than those in PrePPI. PrePPI predictions for non-direct interactions also provide valuable information by identifying pairs of proteins that might be present in multi-protein complexes and, moreover, PrePPI predictions can be used to identify all proteins that are in physical contact in such a complex [2]. PrePPI predictions can also be used in the construction of PPI networks that comprise both direct and indirect interactions and, when combined with features based on context-specific gene expression or knockout screens, can provide insight into dysregulation of cellular signaling as demonstrated with the KRAS-centered OncoSig network for LUAD [23].

Given the continuous developments in structure determination and sequence analysis, PrePPI will continue to evolve and to incorporate new technologies. One possibility is to leverage the proteome-wide, complementary approaches of PrePPI and D-SCRIPT [19] and integrate the interface predictions from both as features in an enhanced PPI prediction algorithm. More computationally intensive methods such as ECLAIR [25] can be used to filter PrePPI predictions thus improving their accuracy. While such methodological advances are contemplated and potentially implemented, the current version of PrePPI will be applied to multiple proteomes and to cross-species interactions as implemented in our P-HIPSTer database [22]. In summary, we believe that PrePPI constitutes a unique resource that will continue to find applications in multiple areas of biomedical science.

## Methods

### Training the SM score

#### Extracting biological interfaces from the PDB

All possible PDB complexes, regardless of source organism, are considered. The quaternary structure of a PDB file frequently does not represent the biologically relevant quaternary structure [26] but will be represented by one of the “biological assemblies” contained in the PDB file. The biological assemblies are specified in the “REMARK 350” lines of the PDB file and contain a set of geometric transformations (“BIOMT” records). A given biological assembly is constructed by applying the transformations defined for that assembly to the set of chains in the PDB file. To define template interface contacts, we construct three-dimensional models of each biological assembly using the associated transformations. A contact between any pair of chains in a biological assembly occurs when two residues from the first chain and two from the second are within 6 Å of each other. The union of these contacts from all biological assemblies for each pair of chains defines the interface for those chains and is used to evaluate structure-based predictions as described in the following sections. ~200K PDB structures, each of which contain, on average, several bioassemblies, are used to construct interfaces.

#### Model Building

Sequences for the human and *E. coli* K12 proteomes are taken from the UniProt defined reference proteomes with one representative protein per gene (Proteome IDs UP000005640 and UP000000625, respectively) [27]. As we recently described [28], each full-length sequence is broken up into individual domains corresponding to those defined in the CDD [21]. Three-dimensional models for each full-length protein are taken from the AlphaFold Protein Structure Database [13] with models for individual domains extracted from the model of the full-length protein. This generates a models database for 75,643 domains from 19,797 human proteins corresponding to 95,440 distinct structural data files and about 4.55 billion pairs for which an interaction is evaluated. Additionally 12,120 domain and full-length sequences for 4,285 *E. coli* proteins.

#### Interaction Model Construction

Sequences for every protein chain in the PDB are downloaded from the PDB web site [3]. The sequences are clustered at a sequence identity cutoff of 60% using the program CD-HIT [29] to form PDB sequence clusters, and a representative for each cluster is defined as the longest sequence in the cluster. For a given query protein, the sequences for its associated models are matched to PDB sequence clusters and then structurally aligned to the PDB structure for the representative of the corresponding cluster. The quality of the structure alignment is scored using the Protein Structural Distance (PSD) calculated using the program ska [30]. Of note, in practice, ska alignments involve protein structures with at least three secondary structure elements so that, beyond PrePPI’s use of sequence orthology as an evidence source, PrePPI typically does not predict interactions involving a single α- helix to a structured domain. If a query model aligns with a PSD<0.6 to the structures of representative sequence of the PDB cluster, the query model is further aligned to all of the member structures. PDB structures with PSD<0.6 are kept as structural neighbors of the query model. Whenever the structures or structural neighbors of two query proteins appear together in a PDB complex (as defined above), we call this complex a “template” for an interaction of the query proteins. In practice, we never create a threedimensional interaction model, rather the structure-based sequence alignments are used to derive properties of the interaction: the quality of the alignment itself; the extent that residues of the query proteins align to interfacial residues in the template; and the extent to which residues predicted to be interfacial in the query proteins align to interfacial residues in the template [1]. Predicted interfacial residues are obtained from our program PredUs 2.0 [14]. This scoring avoids the need to explicitly calculate pairwise properties while preserving context-specific information for the template complex and enables rapid evaluation of interaction models from among billions of possible pairwise query combinations.

Given that the full length protein and multiple domains are used for each protein and multiple models are tested for each of the 95K human query sequences, 10s of billions of interaction models must be evaluated. Each model is evaluated using a scoring function derived from a Bayesian network based on features as summarized above and reported previously [2]. Training of the Bayesian network is based on training sets as described below. For a given protein pair, the highest scoring interaction, whether it is between two full length proteins or between two domains, is chosen for that PPI, leading to about 200 million scored predictions.

#### True positive data sets

The most obvious training set for direct interactions is the PDB [3] it contains a relatively limited number of entries for complexes in a given proteome and redundancies further limit this number. Instead, we have preferred to use the HINT high-quality literature-curated database, HINT HQ-LC [10], which appears to be the best source for direct physical interactions and currently has 16K entries for human and 1,753 for *E. coli*.

We have used a number of databases to calculate ROC curves. The size of these databses and the overlap between them appears in Table 1. They include:

##### Interactome3D [16]

PDB structures and easily constructed homology models.

##### HINT high-quality literature-curated (HINT HQ-LC) [10]

Binary PPIs with at least two literature references.

##### APID Level 2 [9]

Interactions proven by at least 1 binary method.

##### STRING-Physical [8]

Direct and indirect PPIs in the same complex with experimental evidence.

##### BioGRID-MV [18]

PPIs curated from both high-throughput datasets and individual focused studies that are validated by multiple experiments.

##### HURI [7]

Binary PPIs validated by three variations of the Y2H assay.

Overall, the lack of overlap among different databases highlights questions about how they are used/chosen in the training of computational methods, especially for those focused on direct interactions. Our decision to train structural clues on a different true positive set than used for non-structural clues is an attempt to address this issue. Both for human and *E. coli*, HINT HQ-LC has significant overlap with Interactome3D consistent with its focus on direct interactions.

#### True negative data set

The negative set used in training and testing consists of all possible human PPIs minus the union of PPIs that appear at any level of confidence in the databases listed in the previous section. The treatment of every interaction for which there is no evidence as a true negative obviously diminishes apparent performance. But our experience has been that, as opposed to precision/recall curves, ROC curves are not significantly affected by the size of the negative set. We have confirmed this behavior by changing the size of the negative set to be 10 times the size of the positive set and found that this has essentially no effect on the various ROC curve statistics (data not shown).

### Training non-structural clues

As reported previously, in addition to structural evidence, PrePPI uses a number of non-structural clues including partner redundancy, GO annotation, sequence orthology, and phylogenetic profile. Details about the calculation and training of non-structural clues are described in our 2016 publication [2] and will not be repeated here. Briefly, the true positive set was taken from multiple databases with the requirement that a PPI be identified in two independent literature references and no attempt was made to distinguish direct physical from non-direct interactions.

## Acknowledgments

This work was supported by the National Institute of Health (grant R35-GM139585).

## Notes

https://honiglab.c2b2.columbia.edu/PrePPI

### Competing Interest Statement

The authors have declared no competing interest.

